# Driving the expression of the *Salmonella enterica* sv Typhimurium flagellum using *flhDC* from *Escherichia coli* results in key regulatory and cellular differences

**DOI:** 10.1101/333146

**Authors:** Ayman Albanna, Martin Sim, Paul A Hoskisson, Colin Gillespie, Christopher V. Rao, Phillip Aldridge

## Abstract

The flagellar systems of *Escherichia coli* and *Salmonella enterica* exhibit a significant level of genetic and functional synteny. Both systems are controlled by the flagellar specific master regulator FlhD_4_C_2_. Since the early days of genetic analyses of flagellar systems it has been known that *E. coli flhDC* can complement a Δ*flhDC* mutant in *S. enterica*. The genomic revolution has identified how genetic changes to transcription factors and / or DNA binding sites can impact the phenotypic outcome across related species. We were therefore interested in asking: using modern tools to interrogate flagellar gene expression and assembly, what would the impact be of replacing the *flhDC* coding sequences in *S. enterica* for the *E. coli* genes at the *flhDC S. entercia* chromosomal locus? We show that even though all strains created are motile, flagellar gene expression is measurably lower when *flhDC*_EC_ are present. These changes can be attributed to the impact of FlhD_4_C_2_ DNA recognition and the protein-protein interactions required to generate a stable FlhD_4_C_2_ complex. Furthermore, our data suggests that in *E. coli* the internal flagellar FliT regulatory feedback loop has a marked difference with respect to output of the flagellar systems. We argue due diligence is required in making assumptions based on heterologous expression of regulators and that even systems showing significant synteny may not behave in exactly the same manner.

## IMPORTANCE

The bacterial motility organelle known as the flagellum is shared across many bacterial species. *Escherichia coli* and *Salmonella enterica* have underpinned our appreciation of how bacteria express and assemble the bacterial flagellum for over half a century. We show that even though the *E. coli* and *S. enterica* flagellar systems look genetically identical, they input regulatory signals into the flagellar system differently. Our conclusions are based on experiments where we carefully transfer the master flagellar regulator from *E. coli* into the *S. enterica* chromosome and measure a range of outputs relating to flagellar gene expression, assembly and functional output.

## INTRODUCTION

The flagellum in the enteric bacteria, *Escherichia coli* and *Salmonella enterica*, has been studied extensively for over fifty years and provides the canonical example for bacterial motility. These studies have revealed not only the complex structure of the enteric flagellum but also its role in host colonization, pathogenesis, and cellular physiology (1-4). In addition, these studies have identified many of the complex regulatory processes that coordinate the assembly and control of this exquisitely complex biological machine (3-5).

The flagellum in *E. coli* and *S. enterica* are structurally very similar and are often tacitly assumed to be effectively identical aside from differences in the filament structure. However, in the case of regulation, these assumptions are based more on sequence similarity rather than on actual experimental data (5) (6). Indeed, a number of studies have shown that these two systems are regulated in entirely different manners in response to environmental signals despite strong gene synteny. For example, many common *E. coli* strains are motile only during growth in nutrient-poor conditions whereas many common *S. enterica* strains are motile only during growth in nutrient-rich conditions (7). In addition, *E. coli* is more motile at 30°C than at 37°C whereas motility *S. enterica* is generally insensitive to these temperature differences (8). *E. coli flhDC* are transcribed from a single transcriptional start site that is responsive to OmpR, RcsB and CRP regulation, to name only a few regulatory inputs (8). In contrast *S. enterica flhDC* transcription is significantly more complex with up to 5 transcriptional start sites, albeit with only a subset being responsible for the majority of *flhDC* transcription (9).

Part of the problem is that different questions have been asked when studying the regulation of motility in these two bacterial species. Most studies in *E. coli* have focused on the environmental signals and associate regulatory process that induce bacterial motility. In particular, they have focused on the processes that regulate the expression of the master flagellar regulator, FlhD_4_C_2_ (8). Most studies in *S. enterica*, on the other hand, have focused on the regulatory processes that coordinate the assembly process following induction (4). In particular, they have focused on the downstream regulatory processes induced by FlhD_4_C_2_ (3).

Despite differences in regulation, the protein subunits of master flagellar regulators, FlhC and FlhD, exhibit high sequence similarity sharing 94 and 92% identity, respectively, between *E. coli* and *S. enterica*. Given that modifications to transcription factors and/or promoter structure can lead to divergence in regulatory circuits (10), we were interested in how FlhD_4_C_2_ functions in different genetic backgrounds? Previously, it was shown that *E. coli flhDC* can complement a Δ*flhDC* mutant in *S. enterica*, suggesting that these proteins are functions identical in the two bacterial species (11). However, it is not clear whether they are regulated in the same manner. We, therefore, investigated the impact of replacing the native master regulator in *S. enterica* with the one from *E. coli*. Defining the impact of known FlhD_4_C_2_ regulators such as ClpP, YdiV, FliT and FliZ on the two complexes suggest that these two species have adapted in how they perceive FlhD_4_C_2_. We argue that these phenotypic differences arise from adaptations *E. coli* and *S. enterica* have made during evolution to expand or modify cellular function with respect to movement within specific environmental niches.

## RESULTS

### *Orthologous* flhDC *from* E. coli *can functionally complement* flhDC *in* S. enterica

Given the similarities between the flagellar systems in *S. enterica* and *E. coli*, we sought to determine whether the FlhD_4_C_2_ master regulator is functionally equivalent in these two species of bacteria. To test this hypothesis, we replaced the *flhDC* genes in *S. enterica flhDC*_SE_) with the *flhDC* genes from *E. coli* (*flhDC*_EC_). The reason that we performed these experiments in *S. enterica* rather than *E. coli* was that the flagellar system is better characterized in the former, particularly with regards to transcriptional regulation. To avoid plasmid associated artefacts associated with the ectopic expression of *flhDC,* we replaced the entire *S. enterica flhDC* operon with the *flhDC* operon from *E. coli* at the native chromosomal locus (Figure S1).

We first tested whether *flhDC*_EC_ was motile as determined using soft-agar motility plates. As shown in Figures 1A and B, these strains formed rings similar to the wild type. These results demonstrate that *flhDC*_EC_ is functional in *S. enterica.* However, motility plates measure both motility and chemotaxis and do not provide any insights regarding possibly changes in the number of flagella per cell. To determine the impact *flhDC*_EC_ had upon flagellar numbers we used a FliM-GFP fusion as a proxy for flagellar numbers (Figure 1C). When this fluorescent protein fusion is expressed in cells, it forms spots associated with nascent C-rings that loosely correlate with the number of flagella (12-14). By counting the number of spots per cell, we can determine the number of flagella made per cell. As shown Figure 1C, *flhDC*_EC_ did not change flagellar numbers as compared to the wild type. These results demonstrate *flhDC*_EC_ induces flagellar gene expression at similar levels as the wild type.

**Figure 1.**
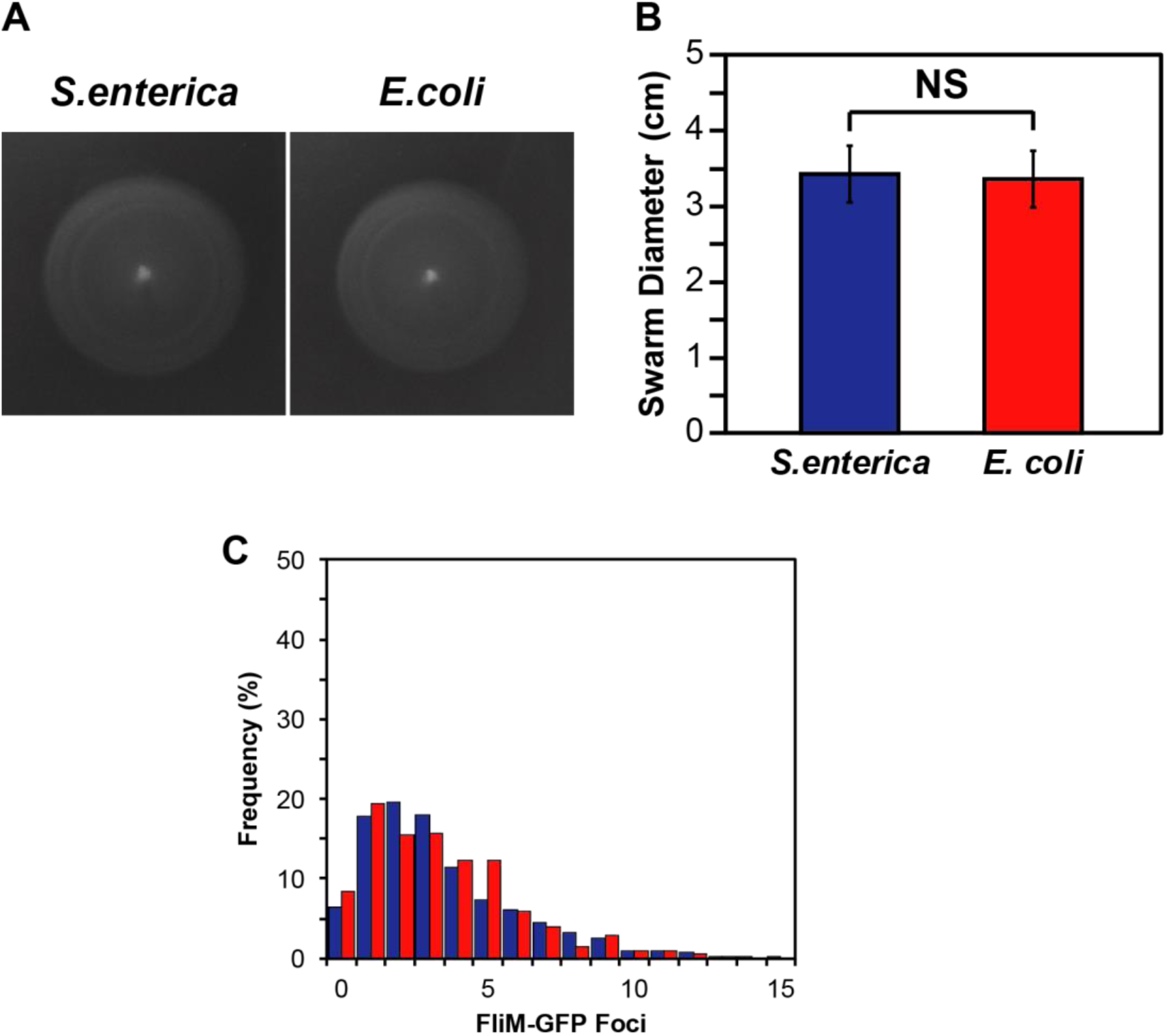
**A.** Motility of *flhDC*_*ST*_ and *flhDC*_*EC*_ driven by P_*flhDC*_. **B.** Quantification of swarms produced in motility agar after 6 to 8 hours incubation. Error bars indicate calculated standard deviations. **C.** Percentage frequency of FliM-GFP foci for *flhDC*_EC_ compared to *S. enterica* with *flhDC* under the control of P_*flhDC*_. Colors of bars in the graph correspond to the source of *flhDC* as shown in (**B**).

### flhDC requires a specific transcription rate to maintain optimal flagellar numbers

The flagellar network in *S. enterica* contains a number of feedback loops to ensure that the cells regulate the number of flagella produced (4). One possibility is that these feedback loops mask any differences in FlhD_4_C_2EC_ activity. To test this hypothesis, we replaced the native P_*flhD*_ promoter with the tetracycline-inducble P_*tetA*_/_*tetR*_ promoters. We then measured flagellar gene expression using a luciferase reporter system (15). In this case, a consistent and significant change in flagellar gene expression was observed when comparing FlhD_4_C_2EC_ to FlhD_4_C_2SE_ activity (Figure 2). Maximal expression of P_*flgA*_ and P_*fliC*_, chosen to reflect flagellar gene expression at different stages of flagellar assembly (5), for both complexes was observed between 10 and 25 ng/ml of anhydrotetracycline, when *flhDC* transcription was from P_*tetA*_ (Figure 2A and B). In contrast, P_*tetR*_, the weaker of the two tetracycline inducible promoters, reached a maximal output between 50 to 100 ng/ml anhydrotetracycline. In both scenarios the output for FlhD_4_C_2EC_ control was lower than for the native FlhD_4_C_2SE_ complex (Figure 2A and 2B).

**Figure 2.**
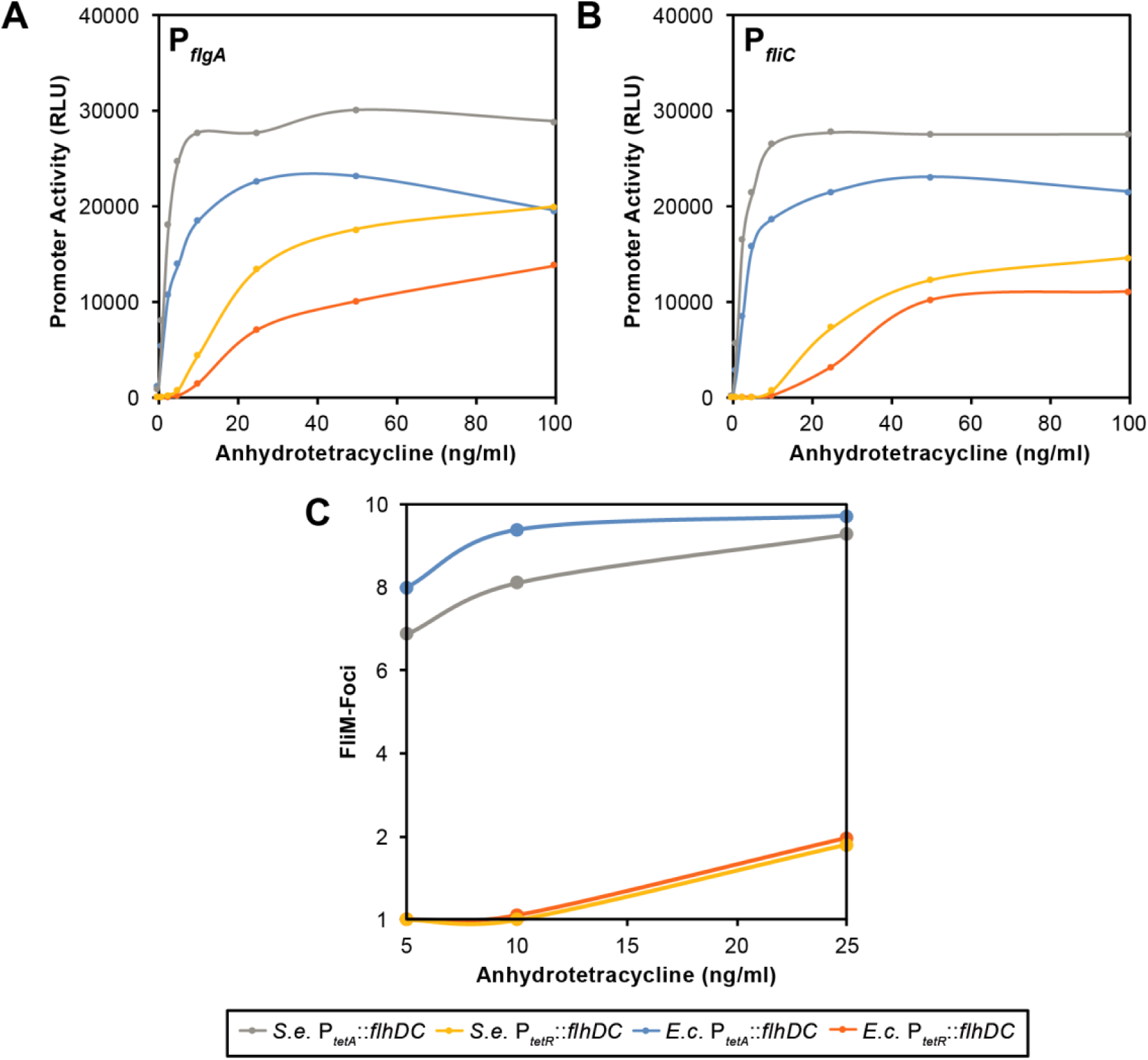
Titration of P_*tetA*_*::flhDC*_*ST/EC*_ and P_*tetR*_*::flhDC*_*ST/EC*_ activity suggests a given rate of transcription drives optimal flagellar assembly. **A.** Activity of P_*flgA*_ in response to P_*tetA*_ or P_*tetR*_ transcription of *flhDC* from *S. enterica* (S.e.) or *E. coli* (E.c.). **B.** Activity of P_*fliC*_ in response to P_*tetA*_ or P_*tetR*_ transcription of *flhDC*. **C.** flagellar numbers as defined by FliM-foci in response to P_*tetA*_ or P_*tetR*_ transcription of *flhDC.*

We also measured the number of FliM-GFP foci at different anhydrotetracycline concentrations. P_*tetR*_::*flhDC* expression generated on average of approximately two FliM-foci per cell at 25 ng/ml of anhydrotetracycline for both FlhD_4_C_2_ complexes (Figure 2C). In contrast, 5 ng/ml induction of the P_*tetA*_::*flhDC*_*EC*_ strain was sufficient to generate typical FliM-foci numbers (approx. 8 flagellar foci per cell). Even with the strong decrease in average foci per cell at these levels of induction, the number of basal bodies observed is sufficient to allow motility at comparable levels in the motility agar assay (Figure S2).

### *Replacement of* flhC *but not* flhD *in* S. enterica *with the* E. coli *orthologs affects motility*

The hetero-oligomeric regulator FlhD_4_C_2_ is unusual in bacteria as the majority of transcriptional regulators are believed to be homo-oligomeric complexes. To determine the relative contributions of the two subunits, we individually replaced the *flhC* or *flhD* genes from *S. enterica* with their ortholog from *E. coli* (Figure S1). When we tested the two strains using motility plates, we found that motility was inhibited in the strain where *flhC*_EC_ replaced the native *S. enterica flhC* (Figure 3A; blue bars), with an 88% reduction in swarm diameter when compared to WT *S. enterica*. The introduction of *flhD*_EC_ compared to *flhDC*_EC_ or *flhDC*_SE_ produced swarms of a comparable size (Figure 3A; blue bars).

**Figure 3.**
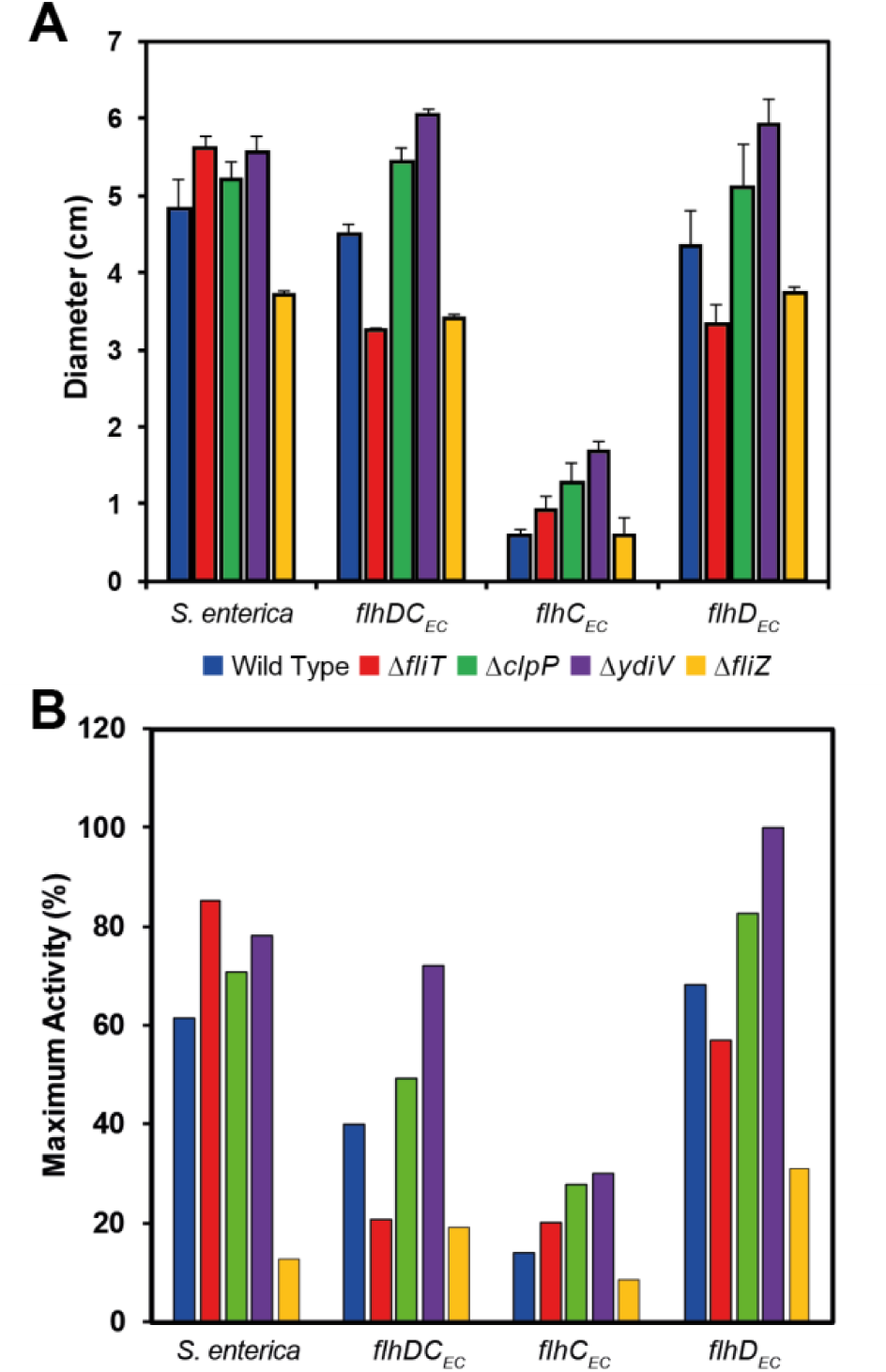
Motility phenotypes and gene expression of *flhDC*_ST_, *flhDC*_EC_, *flhD*_EC_ and *flhC*_EC_ strains in the absence of known FlhD_4_C_2_ regulators. **A.** Quantification of n = 3 swarms per strain produced in motility agar after 6 to 8 hours incubation at 37°C. Error bars indicate calculated standard deviations. **B.** Relative activity of P_*fliC*_ in all strains as a percent of the maximal activity observed in *flhD*_EC_ Δ*ydiV*.

Using the dose-dependent inducible P_*tetA*_ promoter(16) we observed that P_*tetA*_ expression of *flhC*_EC_ led to reduced P_*flgA*_ transcription and strongly reduced P_*fliC*_ transcription (Figure 4). Strains expressing *flhD*_EC_ in *S. enterica* showed a mild increase in P_*flgA*_ gene expression and a similar response for P_*fliC*_, although these changes were not significant (P = 0.32) (Figure 4). These data suggest that the combination of FlhD_SE_ and FlhC_EC_ generates an inefficient FlhD_4_C_2_ complex, resulting in reduced motility.

**Figure 4.**
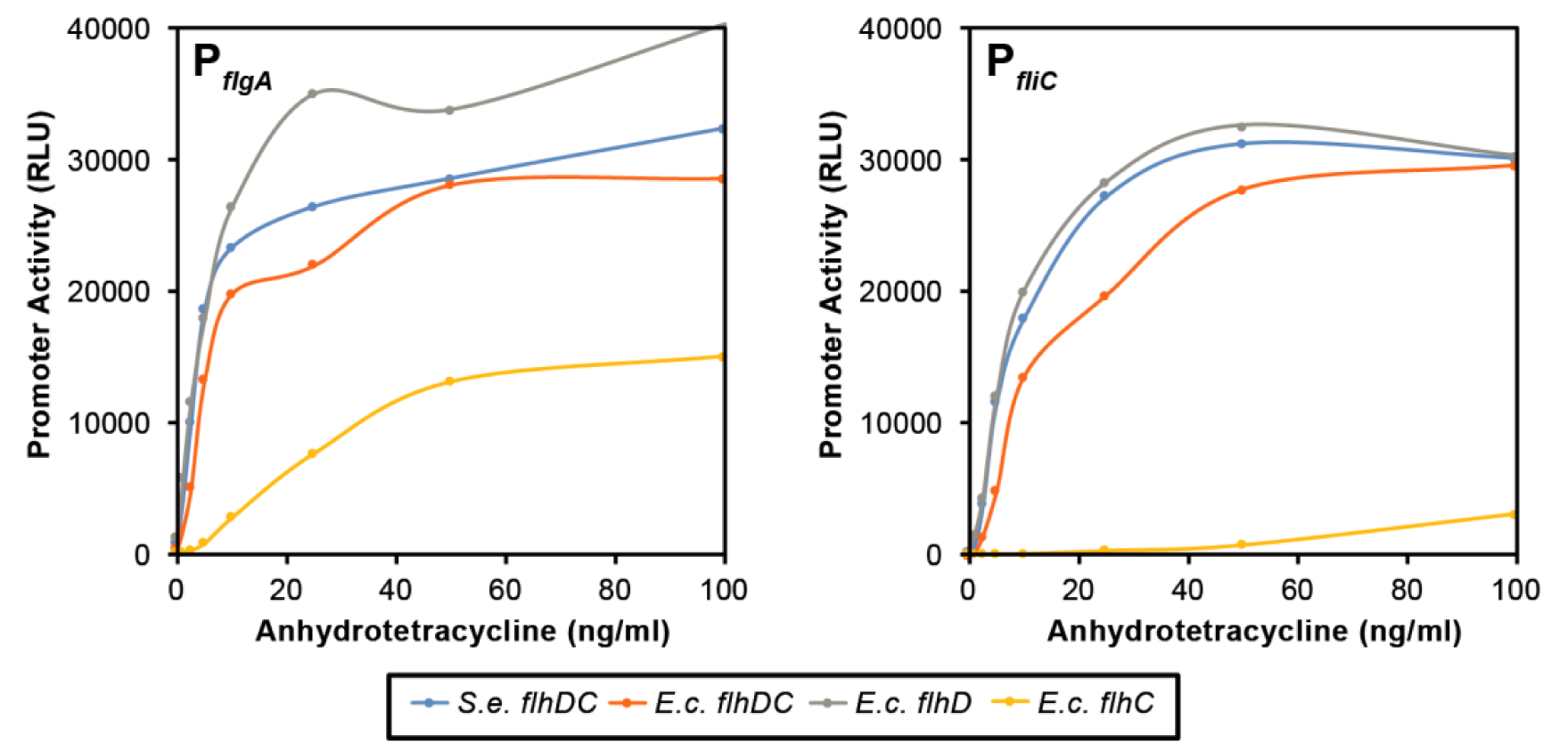
Titration of P_*tetA*_*::flhDC* for *S. enterica*, *flhDC*_EC_, *flhD*_EC_ and *flhC*_EC_ suggests that *flhC*_EC_ exhibits low motility due reduced P_*flgA*_ activity and a strong reduction in P_*fliC*_ activity. Inducible expression was driven from the P_*tetA*_ promoter within the TetRA cassette of Tn10. The data shown in both panels is significant using ANOVA statistical analysis P < 0.05.

### Orthologous FlhC and FlhD interaction is species specific and a key determinant of promoter recognition by the FlhD_4_C_2_ complex

The results above demonstrate that *flhC*_EC_ is not functionally identical to *flhC*_ST_. One possibility is that that FlhC_EC_ is impaired in FlhD_4_C_2_ for DNA-binding. Alternatively, the stability of the FlhD_4_C_2_ complex is reduced in the *flhC*_*EC*_ strain, leading to reduced FlhD_4_C_2_ activity. To test these hypotheses, we purified all combinations of the FlhD_4_C_2_ complex using affinity (Ni+ and heparin) chromatography (Figure 5A). In each complex, FlhD was tagged with a carboxy-terminal hexa-histidine to facilitate affinity purification. Such expression constructs have previously been used successfully to purify the FlhD_4_C_2_ complex (17, 18). Using either Ni+ affinity or heparin purification, we observed complete complex retrieval for three combinations (Figure 5A). FlhC recovery was less efficient in the FlhD_SE_/FlhC_EC_ complex. In contrast, no FlhD_SE_/FlhC_EC_ complex was recovered via Heparin purification, used to mimic DNA during protein purification of DNA-binding proteins (Figure 5A). This suggests that the FlhD_SE_/FlhC_EC_ complex is less stable, resulting on a lower yield of complex retrieval.

**Figure 5.**
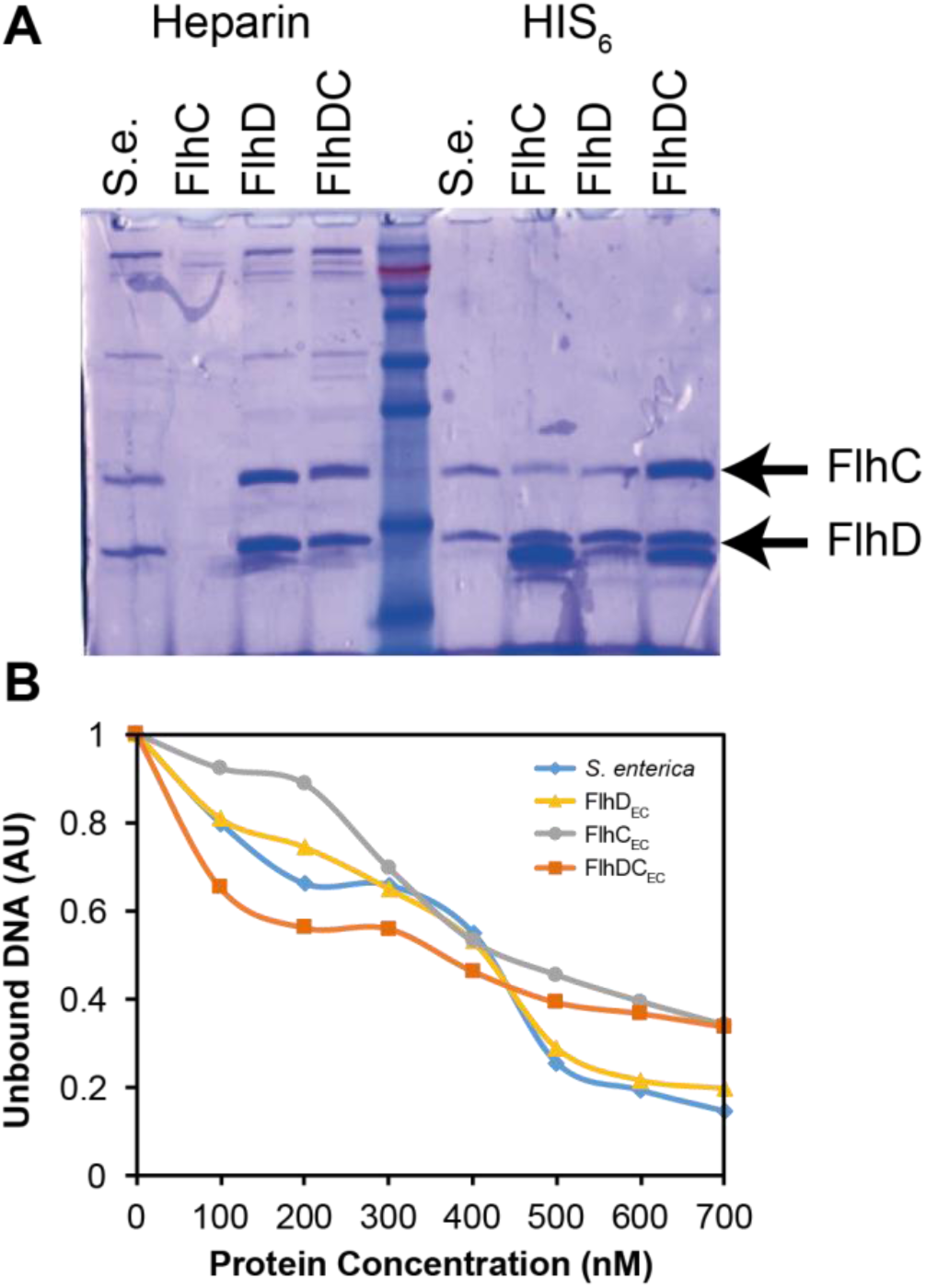
The FlhD_ST_FlhC_EC_ complex is an active but unstable complex. **A.** Protein gel showing purified complexes with either HIS_6_ or Heparin based purification protocols. The nature of the FlhDC complex allows isolation of both proteins in these assays. Arrows indicate the FlhC and FlhD bands. **B.** Quantification of EMSA to define the binding ability of the complex combinations compared to *S. enterica* FlhD_4_C_2_. Data shows that whenever FlhC_EC_ is present a reduced level of binding to P_*flgAB*_ was observed.

We next used the EMSA assays to test all four protein complexes for their ability to bind the *S. enterica* P_*flgAB*_ promoter region. Quantification of the DNA shifts showed that complexes containing the orthologous FlhC_EC_ reduced the P_*flgAB*_ promoter binding profile, compared to FlhC_SE_ complexes (Figure 5B). This is consistent with FlhC being the DNA binding subunit of the complex and the variation in FlhD_4_C_2_ activated promoter-binding sites between *S. enterica* and *E. coli* (19). Therefore, these results suggest that FlhC is a key determinant of DNA binding ability. Furthermore, the reduction in FlhC_EC_ motility and flagellar gene expression in *S. enterica* is a result of the FlhD_SE_/FlhC_EC_ complex being unstable, ultimately reducing the cellular concentration of the FlhD_4_C_2_ complex.

### FlhD_4_C_2EC_ responds to proteolytic regulation

*S. enterica* and *E. coli* both regulate the FlhD_4_C_2_ complex through ClpXP-mediated proteolytic degradation. Proteolytic degradation of FlhD_4_C_2_ plays a fundamental role in facilitating rapid responses to environmental changes that require motility (20, 21). The FlhD_4_C_2_ complex has a very short half-live of approximately 2-3 minutes (22). Proteolytic degradation of FlhD and FlhC is regulated in *E. coli* and *S. enterica* by YdiV (23).

However, *ydiV* is not expressed under standard laboratory conditions in model *E. coli* strains, suggesting that ClpXP activity is modulated in a species-specific manner (7).

Previous work has shown that YdiV delivers FlhD_4_C_2_ complexes to ClpXP for degradation (24). We have assessed the impact on motility for Δ*clpP* and Δ*ydiV* mutations (Figure 3). The Δ*clpP* and Δ*ydiV* mutants exhibited improved motility and flagellar gene expression, including the FlhD_SE_/FlhC_EC_ strain (Figure 3A and B). These results suggest that proteolytic degradation mechanism of FlhD and FlhC, and its regulation, is common to *E. coli* and *S. enterica*.

To complement the motility assays, we investigated how Δ*clpP* and Δ*ydiV* mutations impact the number of FliM-foci in cell. Both Δ*clpP* and Δ*ydiV* mutants showed an increased number of FliM-foci compared to the wild type (Figure 6 A-C). For *flhC*_EC_ strain, FliM-foci were observed in 13% of the population where individual cells exhibited just one or two foci. However, the Δ*clpP* or Δ*ydiV* mutants increased the flagellated population of the *flhC*_EC_ strains to 51 and 46 % respectively, albeit with the majority still possessing only a single FliM focus (Figure 6 B and C).

**Figure 6.**
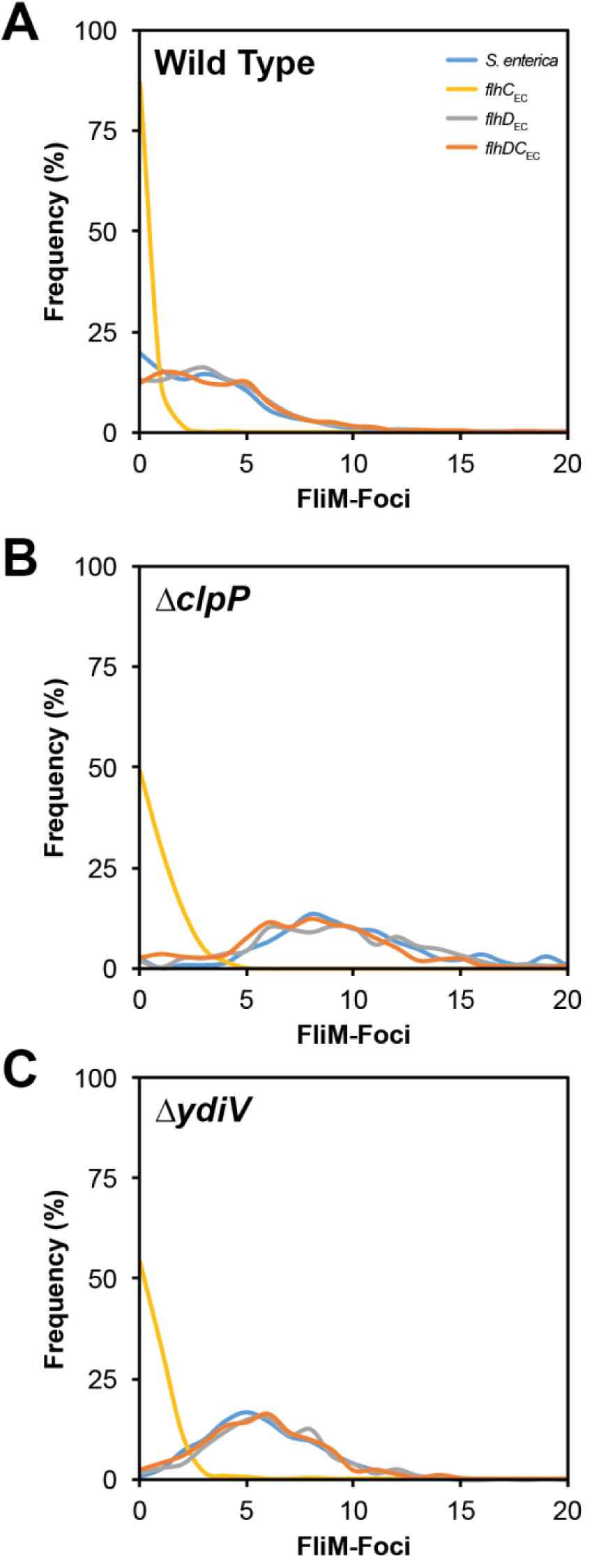
Impact of protein stability regulators of FlhD_4_C_2_ on flagellar numbers as defined by FliM-foci. Quantification of FliM-foci was performed using the semi-automatic protocols defined with in Microbetracker. **A.** Wild Type foci distribution; **B.** Δ*clpP*; **C.** Δ*ydiV*.

### FliT and FliZ regulation of FlhD_4_C_2_ complexes

FlhD_4_C_2_ activity has an additional level of regulation in *S. enterica* via the flagellar-specific regulators FliT and FliZ. FliT functions as an export chaperone for the filament cap protein, FliD, and is a regulator of FlhD_4_C_2_ activity (17, 25). FliT disrupts the FlhD_4_C_2_ complex but is unable to disrupt a FlhD_4_C_2_:DNA complex. Therefore, FliT modulates availability of FlhD_4_C_2_ complexes for promoter binding (17). In contrast, FliZ is a negative regulator of *ydiV* expression and thus increases the number of FlhD_4_C_2_ complexes in *S. enterica* (26, 27).

In motility assays of Δ*fliT* mutants, we observed a difference between our different *flhDC* strains. Motility is increased in a Δ*fliT* mutant background in *S. enterica* ((28) and Figure 3A). However, when *flhDC*_EC_ and *flhD*_EC_ replaced the native genes, a reduced swarm size was observed (Figure 3A). Furthermore, quantification of P_*fliC*_ activity agreed with the motility profile for Δ*fliT* mutants, where *flhDC*_EC_ and *flhD*_EC_ containing strains had reduced promoter activity compared to wild type (Figure 3B). This suggests that the FlhD_4_C_2_ complexes are being perceived differently by FliT in *S. enterica*. The results for Δ*clpP* and Δ*ydiV* mutants suggests that this is not due to protein stability, as all complex combinations reacted in a comparable fashion (Figure 3).

In contrast, the loss of *fliZ* resulted in a consistent reduction in motility, except for the *flhC*_EC_ strain. However, as the *flhC*_EC_ strain was already impaired in motility, it is possible that the resolution of the motility assay was unable to identify differences in Δ*fliZ* mutant. Flagellar gene expression activity did, however, suggest a 2-fold drop in P_*fliC*_ expression in the *flhC*_EC_ Δ*fliZ* strain as compared to the otherwise wild-type (Figure 3B).

Analysis of FliM-foci distribution in Δ*fliT* mutant reinforced the observed discrimination of *flhDC*_EC_ and *flhD*_EC_ gene replacements. Calculating the average foci per cell, *S. enterica* Δ*fliT* mutants showed an increased average number of foci per cell from 2.9 to 6.3, while the *flhD*_EC_ (*fliT*^+^: 3.4 versus Δ*fliT*: 4.2) and *flhDC*_EC_ replacements (*fliT*^+^: 3.6 versus Δ*fliT*: 2.7) exhibited no significant changes (Figure 7A). Interestingly, in a Δ*fliZ* mutant background, the FliM-foci analysis was able to differentiate *flhDC*_EC_ and *flhD*_EC_ from the native *S. enterica flhDC* strain. Both replacements exhibited an increase in the average foci compared to *S. enterica* Δ*fliZ* (Figure 7A).

**Figure 7.**
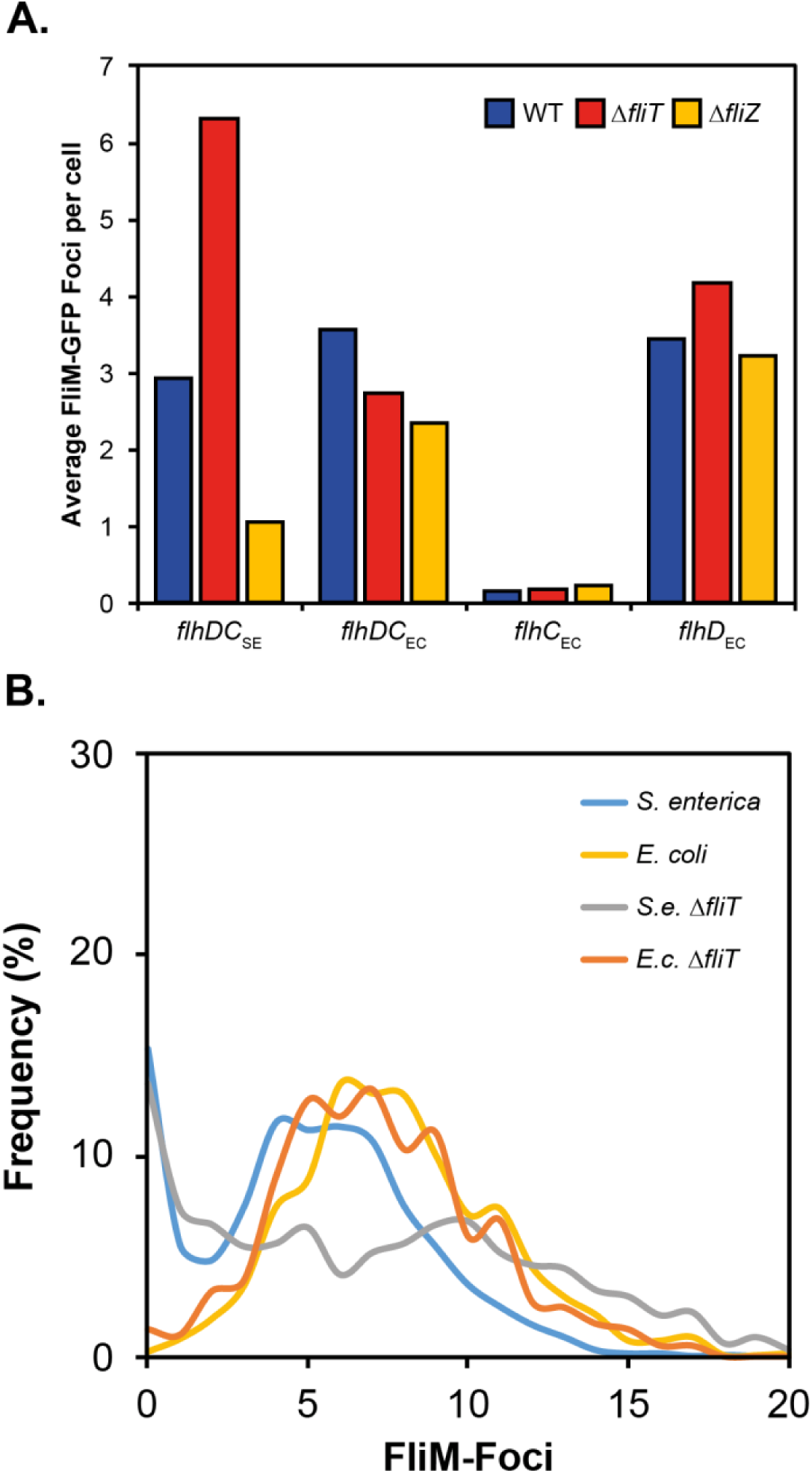
FliT and FliZ regulation reflects when FlhC_EC_ or FlhD_EC_ are present. **A.** FliM-Foci quantification is consistent with the observed motility phenotype of Δ*fliT* mutants. For Δ*fliZ* FliM-foci numbers discriminate between the source of FlhD, FlhD_SE_ exhibits a consistnet drop in foci while FlhD_EC_ containing strains show comparable foci averages. **B.** Testing the hypothesis that Δ*fliT* mutants respond differently in *E. coli* compared to *S. enterica*. Note: this experiment in **(B)** uses the species *E. coli* and *S. enterica* not engineered replacements.

These data suggest that there is a fundamental difference in how the FlhD_4_C_2_ complexes in *E. coli* and *S. enterica* respond to, at least, FliT regulation. There are two explanations for this: a) the *E. coli* combinations are being regulated via an unidentified mechanism in *S. enterica* or b) that they are insensitive to FliT regulation. Both arguments predict that in *E. coli* FlhD_4_C_2_ may respond differently to FliT regulation. Comparing *S. enterica* and *E. coli* does indeed identify a difference in the response to a Δ*fliT* mutant. While a Δ*fliT* mutant in *S. enterica* leads to a consistent increase in FliM-foci, no significant difference is noted for an *E. coli* Δ*fliT* mutant compared to *E. coli* wild type (Figure 7B). This suggests that the regulatory impact of FliT is very different in these two flagellar systems and the role FliT plays in *S. enterica* is potentially adaptive and species specific.

## DISCUSSION

Two model flagellar systems that form the foundation of the flagellar field are those from the enteric species *E. coli* and *S. enterica*. These two systems have led to key discoveries in relation to many aspects of flagellar structure, type 3 secretion, flagellar cell biology and the regulation of flagellar assembly. Textbook explanations suggest that most flagellar systems are being activated, regulated and built according to the models for *E. coli* and *S. enterica*. Modifications of transcriptional regulatory circuits contribute to the phenotypic diversity we see in closely related gene sets and we are only now able to investigate this in depth due to the tools available. Here we have taken a simple step and asked how do orthologous FlhD_4_C_2_ complexes function in the closely related species *E. coli* and *S. enterica?*

At the onset of our work it was known that FlhD_4_C_2_ from *E. coli* could sustain motility in *S. enterica*(11). Our work was focussed on understanding and defining the species-specific differences in the regulon of two orthologous genes. Here we took advantage of the well-defined flagellar assembly tools to measure outputs such as, motility, flagellar assembly per cell and flagellar gene expression. Bioinformatic analysis identifies only an 8 and 6% identity difference between FlhD and FlhC in *E. coli* and *S. enterica* respectively, suggesting that these proteins function in an analogous fashion. It is well established that related taxa usually rely on orthologous regulators to coordinate response to a given signal (10).

The fine detail of the differences in the FlhD_4_C_2_ complexes only became apparent when we began to focus on their effect on flagellar gene expression and flagellar assembly. In all of our assays FlhD_4_C_2EC_ exhibited a reduction in flagellar gene expression compared to FlhD_4_C_2SE_. Biochemical analysis of isolated complexes showed that FlhC_EC_ had weaker DNA binding ability to the P_*flgAB*_ promoter region from *S. enterica,* consistent with previous investigations into FlhD_4_C_2_ DNA binding activity (19). The isolation of FlhD_4_C_2_ complexes from our strains suggested that a key aspect of the phenotypes we observed, was the stability of the complexes formed.

With respect to *flhDC* transcription we show a discrepancy in flagellar numbers defined by FliM-foci when using P_*tetA*_/P_*tetR*_::*flhDC* expression. This was somewhat surprising as all constructs exhibited good swarming ability on motility agar plates (Figure S2). Original studies on the regulation of P_*tetA*_/P_*tetR*_ from Tn10 have shown that these two promoters have differing activities but both respond to TetR regulation. We show that even though maximal activity of P_*flgA*_ and P_*fliC*_ can reach 40-50% of P_*tetA*_::*flhDC* expression for P_*tetR*_ strains, this results in an average of 2 flagella per cell. This suggests that even though the majority of the literature states that *E. coli* and *S. enterica* produce between 4 and 8 flagella per cell, only 1 or 2 per cell is needed for an optimal output of the system with respect to motility agar assays.

It has been shown that FliT interacts with FlhC and that in *S. enterica* the output of this circuit is to destabilize FlhD_4_C_2_ complexes that are not bound to DNA. Our data suggests that this level of regulation does not impact *E. coli* FlhC. The nature of the adaptability needed by the favourable conditions to drive motility in *E. coli* may have led to the FliT regulatory input becoming less critical. Similarly, the impact of FliZ regulation becomes apparent for FlhD_EC_ containing complexes when we assess flagellar numbers. FliZ regulates the transcription of *ydiV* in *S. enterica* (27). It is plausible that the impact in changing *ydiV* regulation is the source of this differentiation, especially as YdiV is proposed to interact with FlhD_SE_. Furthermore, we know that *ydiV* is not expressed in model *E. coli* strains, strengthening the argument that FlhD_EC_ has adapted to the absence of YdiV or vice versa FlhD_SE_ to YdiV.

Importantly our analysis shows that even though these two systems are genetically similar, investigation of FlhD_4_C_2_ activity identifies subtle but key differences into how the FlhD_4_C_2_ complex is modulated in two closely related species. We argue that this is a valid example of the caution needed in the age of synthetic biology to exploit heterologous systems in alternative species or chassis’. Our data shows that even systems showing significant synteny may not behave in exactly the same manner and due diligence is required in making assumptions based on heterologous expression.

## MATERIALS AND METHODS

### Bacterial Strains and Growth conditions

*S. enterica* and *E. coli* strains used in this study have been previously described elsewhere (12, 15, 17, 28). This study used *S. enterica* serovar Typhimurium strain LT2 as the chassis for all experiments. *E. coli* genetic material was derived from MG1655. All strains were grown at either 30°C or 37°C in Luria Bertani Broth (LB) either on 1.5% agar plates or shaken in liquid cultures at 160 rpm (17). Antibiotics used in this study have been described elsewhere (29). Motility assays used motility agar (17) incubated at 37°C for 6 to 8 hours. Motility swarms were quantified using images captured on a standard gel doc system with a ruler in the field of view and quantified using ImageJ to measure the vertical and horizontal diameter using the average as the swarm size. All motility assays were performed in triplicate using single batches of motility agar.

### Genetic Manipulations

For the replacement of *flhDC* coding sequences the modified lambda red recombination system described by Blank et al (2011) was used (30). Deletion of *clpP, ydiV, fliT* and *fliZ* was performed using the pKD system described by Datsenko and Wanner (2000) (31).

P_*tetA*_ / P_*tetR*_ replacements of the P_*f lhDC*_ region was also performed using Datsenko and Wanner (2000) with the template being Tn10*d*Tc (32). For Blank et al (2011) replacement experiments we used autoclaved chlortetracycline instead of anhydrotetracycline as described for the preparation of Tetracycline sensitive plates (33). All other gene replacements were performed as previously described (17). All primers used for these genetic manipulations are available on request.

### Quantification of flagellar gene expression

Flagellar gene expression assays were performed using the plasmids pRG39::cat (P_*fliC*_) and pRG52::cat (P_*flgA*_) (15). Both plasmids were transformed into strains using electroporation. Gene expression was quantified as described previously and analysis was based on a minimum of n = 3 repeats for each strain tested (15).

### Quantification of FliM-GFP foci

FliM-GFP foci were quantified using Microbetracker on images captured using a Nikon Ti inverted microscope using filters and exposure times described previously (14). Strains were grown to an OD600 of 0.5 to 0.6 and cells immobilised using a 1 % agarose pad containing 10 % LB (14, 17). For each strain a minimum of 5 fields of view were captured from 3 independent repeats. This allowed analysis of approximately 1000 - 1500 cells per strain. For the comparison of FliM foci in *E. coli* Δ*fliT* to *S. enterica* Δ*fliT* shown in Figure 7B the chemostat growth system described by Sim et al (2017) was used. For this experiment the growth rate of both strains was similar to batch culture in LB at 37°C where the media used was a MinE base with 0.1% Yeast extract and 0.2% glucose (14, 17).

### Purification of FlhD_*4*_*C*_*2*_ complexes

Purification of proteins complexes was based on previously described methods (17). Wild type FlhD_4_C_2SE_ was purified using pPA158. The other 3 complexes were purified from plasmids generated using the New England Biolabs NEBuilder DNA Assembly kit on the backbone of pPA158. The *E. coli* strain BL21 was used for all protein induction experiments prior to protein purification using either a pre-equilibrated 5ml His-trap column or a 5ml heparin column (GE Healthcare). Proteins were visualised using Tricine-based SDS polyacrylamide gel electrophoresis and standard commassie blue staining (17).

### Electrophoretic mobility shift assay (EMSA)

All EMSA assays were performed using Ni++ (his-trap) purified proteins as this allowed analysis of all four complexes (Figure 5A). Buffer exchange from elution buffer to a 100mM Tris-HCl, 300 mM NaCl 1mM DTT (pH 7.9) buffer was performed through 10 cycles of protein concentration in VivaSpin columns with 20 ml buffer reduced to 5 ml per round of centrifugation at 4500 rpm. A protein concentration range of 100 to 700 nM was used with 80 ng / ml of a PCR product containing P_*flgAB*_ from *S. enterica*. After incubation bound and unbound DNA were resolved using 5% acrylamide gels made with 1x TBE buffer.

Quantification of gel images was performed using ImageJ.

## ACKNOWLEDGEMENTS

PDA would like to recognize the internal financial support of ICAMB during this study. The stipend and research costs for the PhD of AA was provided by The Ministry of Higher Education and Scientific Research (Iraq). We would like to thank the financial support of the Newcastle University Faculty of Medicine for providing the John W illiam Luccok and Ernest Jeffcock Research PhD Studentship to MS for this study. PAH would like to acknowledge the support of iUK/BBSRC (grant: BB/N023544/1), NERC (grant: NE/M001415/1), the University of Strathclyde and the Microbiology Society for funding. We would also like to thank all lab members for feedback on the project during the experimental and writing phases.

## REFERENCES

1. Duan Q, Zhou M, Zhu L, Zhu G. 2013. Flagella and bacterial pathogenicity. J Basic Microbiol 53:1–8.

2. Minamino T, Imada K, Namba K. 2008. Mechanisms of type III protein export for bacterial flagellar assembly. Mol Biosyst 4:1105–1115.

3. Chevance FFV, Hughes KT. 2008. Coordinating assembly of a bacterial macromolecular machine. Nat Rev Micro 6:455–465.

4. Aldridge P, Hughes KT. 2002. Regulation of flagellar assembly. Curr Opin Microbiol 5:160–165.

5. Chilcott GS, Hughes KT. 2000. Coupling of flagellar gene expression to flagellar assembly in Salmonella enterica serovar typhimurium and Escherichia coli. Microbiol Mol Biol Rev 64:694–708.

6. Minamino T, Namba K. 2004. Self-assembly and type III protein export of the bacterial flagellum. J Mol Microbiol Biotechnol 7:5–17.

7. Wada T, Hatamoto Y, Kutsukake K. 2012. Functional and expressional analyses of the anti-FlhD4C2 factor gene ydiV in Escherichia coli. Microbiology 158:1533–1542.

8. Soutourina OA, Bertin PN. 2003. Regulation cascade of flagellar expression in Gram-negative bacteria. FEMS Microbiol Rev 27:505–523.

9. Mouslim C, Hughes KT. 2014. The effect of cell growth phase on the regulatory cross-talk between flagellar and Spi1 virulence gene expression. PLoS Pathog 10:e1003987.

10. Perez JC, Groisman EA. 2009. Evolution of transcriptional regulatory circuits in bacteria. Cell 138:233–244.

11. Kutsukake K, Iino T, Komeda Y, Yamaguchi S. 1980. Functional homology of fla genes between Salmonella typhimurium and Escherichia coli. Mol Gen Genet 178:59–67.

12. Aldridge P, Karlinsey JE, Becker E, Chevance FFV, Hughes KT. 2006. Flk prevents premature secretion of the anti-sigma factor FlgM into the periplasm. Mol Microbiol 60:630–643.

13. Delalez NJ, Wadhams GH, Rosser G, Xue Q, Brown MT, Dobbie IM, Berry RM, Leake MC, Armitage JP. 2010. Signal-dependent turnover of the bacterial flagellar switch protein FliM. Proceedings of the National Academy of Sciences 107:11347–11351.

14. Sim M, Koirala S, Picton D, Strahl H, Hoskisson PA, Rao CV, Gillespie CS, Aldridge PD. 2017. Growth rate control of flagellar assembly in Escherichia coli strain RP437. Sci Rep 7:41189.

15. Brown JD, Saini S, Aldridge C, Herbert J, Rao CV, Aldridge PD. 2008. The rate of protein secretion dictates the temporal dynamics of flagellar gene expression. Mol Microbiol 70:924–937.

16. Bertrand KP, Postle K, Wray LV, Reznikoff WS. 1984. Construction of a single-copy promoter vector and its use in analysis of regulation of the transposon Tn10 tetracycline resistance determinant. J Bacteriol 158:910–919.

17. Aldridge C, Poonchareon K, Saini S, Ewen T, Soloyva A, Rao CV, Imada K, Minamino T, Aldridge PD. 2010. The interaction dynamics of a negative feedback loop regulates flagellar number in Salmonella enterica serovar Typhimurium. Mol Microbiol 78:1416–1430.

18. Wang S, Fleming RT, Westbrook EM, Matsumura P, McKay DB. 2006. Structure of the Escherichia coli FlhDC complex, a prokaryotic heteromeric regulator of transcription. Journal of Molecular Biology 355:798–808.

19. Stafford GP, Ogi T, Hughes C. 2005. Binding and transcriptional activation of nonflagellar genes by the Escherichia coli flagellar master regulator FlhD2C2. Microbiology (Reading, Engl) 151:1779–1788.

20. Kitagawa R, Takaya A, Yamamoto T. 2011. Dual regulatory pathways of flagellar gene expression by ClpXP protease in enterohaemorrhagic Escherichia coli. Microbiology 157:3094–3103.

21. Tomoyasu T, Ohkishi T, Ukyo Y, Tokumitsu A, Takaya A, Suzuki M, Sekiya K, Matsui H, Kutsukake K, Yamamoto T. 2002. The ClpXP ATP-dependent protease regulates flagellum synthesis in Salmonella enterica serovar typhimurium. J Bacteriol 184:645–653.

22. Claret L, Hughes C. 2000. Rapid Turnover of FlhD and FlhC, the Flagellar Regulon Transcriptional Activator Proteins, during Proteus Swarming. J Bacteriol 182:833–836.

23. Wada T, Morizane T, Abo T, Tominaga A, Inoue-Tanaka K, Kutsukake K. 2011. EAL domain protein YdiV acts as an anti-FlhD4C2 factor responsible for nutritional control of the flagellar regulon in Salmonella enterica Serovar Typhimurium. J Bacteriol 193:1600–1611.

24. Takaya A, Erhardt M, Karata K, Winterberg K, Yamamoto T, Hughes KT. 2012. YdiV: a dual function protein that targets FlhDC for ClpXP-dependent degradation by promoting release of DNA-bound FlhDC complex. Mol Microbiol 83:1268–1284.

25. Bennett JC, Thomas J, Fraser GM, Hughes C. 2001. Substrate complexes and domain organization of the Salmonella flagellar export chaperones FlgN and FliT. Mol Microbiol 39:781–791.

26. Saini S, Brown JD, Aldridge PD, Rao CV. 2008. FliZ Is a posttranslational activator of FlhD4C2-dependent flagellar gene expression. J Bacteriol 190:4979–4988.

27. Wada T, Tanabe Y, Kutsukake K. 2011. FliZ Acts as a Repressor of the ydiV Gene, Which Encodes an Anti-FlhD4C2 Factor of the Flagellar Regulon in Salmonella enterica Serovar Typhimurium. J Bacteriol 193:5191–5198.

28. Aldridge P, Karlinsey J, Hughes KT. 2003. The type III secretion chaperone FlgN regulates flagellar assembly via a negative feedback loop containing its chaperone substrates FlgK and FlgL. Mol Microbiol 49:1333–1345.

29. Bonifield HR, Hughes KT. 2003. Flagellar phase variation in Salmonella enterica is mediated by a posttranscriptional control mechanism. J Bacteriol 185:3567–3574.

30. Blank K, Hensel M, Gerlach RG. 2011. Rapid and highly efficient method for scarless mutagenesis within the Salmonella enterica chromosome. PLoS ONE 6:e15763.

31. Datsenko KA, Wanner BL. 2000. One-step inactivation of chromosomal genes in Escherichia coli K-12 using PCR products. Proc Natl Acad Sci USA 97:6640–6645.

32. Rappleye CA, Roth JR. 1997. A Tn10 derivative (T-POP) for isolation of insertions with conditional (tetracycline-dependent) phenotypes. J Bacteriol 179:5827–5834.

33. Maloy SR, Nunn WD. 1981. Selection for loss of tetracycline resistance by Escherichia coli. J Bacteriol 145:1110–1111.

